# SARS-CoV-2 emerging complexity

**DOI:** 10.1101/2021.01.27.428384

**Authors:** Francesca Bertacchini, Eleonora Bilotta, Pietro Salvatore Pantano

## Abstract

The novel SARS_CoV-2 virus, prone to variation when interacting with spatially extended ecosystems and within hosts^1^ can be considered a complex dynamic system^2^. Therefore, it behaves creating several space-time manifestations of its dynamics. However, these physical manifestations in nature have not yet been fully disclosed or understood. Here we show 4-3 and 2-D space-time patterns of rate of infected individuals on a global scale, giving quantitative measures of transitions between different dynamical behaviour. By slicing the spatio-temporal patterns, we found manifestations of the virus behaviour such as cluster formation and bifurcations. Furthermore, by analysing the morphogenesis processes by entropy, we have been able to detect the virus phase transitions, typical of adaptive biological systems^3^. Our results for the first time describe the virus patterning behaviour processes all over the world, giving for them quantitative measures. We know that the outcomes of this work are still partial and more advanced analyses of the virus behaviour in nature are necessary. However, we think that the set of methods implemented can provide significant advantages to better analyse the viral behaviour in the approach of system biology^4^, thus expanding knowledge and improving pandemic problem solving.

## Introduction

Months after the official announce of the first case of pneumonia of Sars-Cov-2 in China^5^, the number of cases of Covid-19 is dramatically rising, and is reappearing in countries where it seemed to have dropped. Despite the responses of countries aimed at limiting transmission, the pandemic is still cause millions of deaths. We are dealing with an unprecedented situation and unable to control the viral dynamics^6^. The behaviour of this RNA virus seems to be adaptive and ready to reappear, with progressively cunning evolutionary dynamics^7^ that seem to escape both the countries control policies and traditional forecasting supported by mathematical models^8^, typically used to manage viral behaviour^9^. In the context of viral epidemiology, the hypothesis that viruses behave as complex adaptive systems is gaining ground^10^. However, despite works using the complex systems approach is extensive, there is not yet a global modelling of the evolution of the novel SARS_COV-2 in nature. In this article, we demonstrate that the viral behaviour of the new SARS-COV-2 behaves as a critically organized system^3 4^.

## Results

To detect the viral patterns, a manifold process has been adopted (Fig. 1). We globally modelled infection rates of COVID-19 at a 0.01∘ resolution for each day of the epidemic course by using a Probability Distribution Function (PDF) and a spatial entropy measure applied on this probability distribution. This allowed us to obtain a hypersurface that encompasses space and time, thus obtaining the visualization of the complexity of the pandemic phenomenon, with the opportunity to analyse the virus emerging behaviour. Reducing the dimensionality of the 4D hypersurface, we also obtained a dynamic visualization of the pandemic wave travelling the globe. To accomplish the whole process, we first collected the data, cleaned it up, exactly defining latitude and longitude, and then displayed it on Mercator’s map. Data have been downloaded from the Wolfram Data Repository^11^, considering the different geo-administrative levels as homogeneous, for the 217 Countries that have been involved in the pandemic (Fig 1, 1).

**Fig 1.**
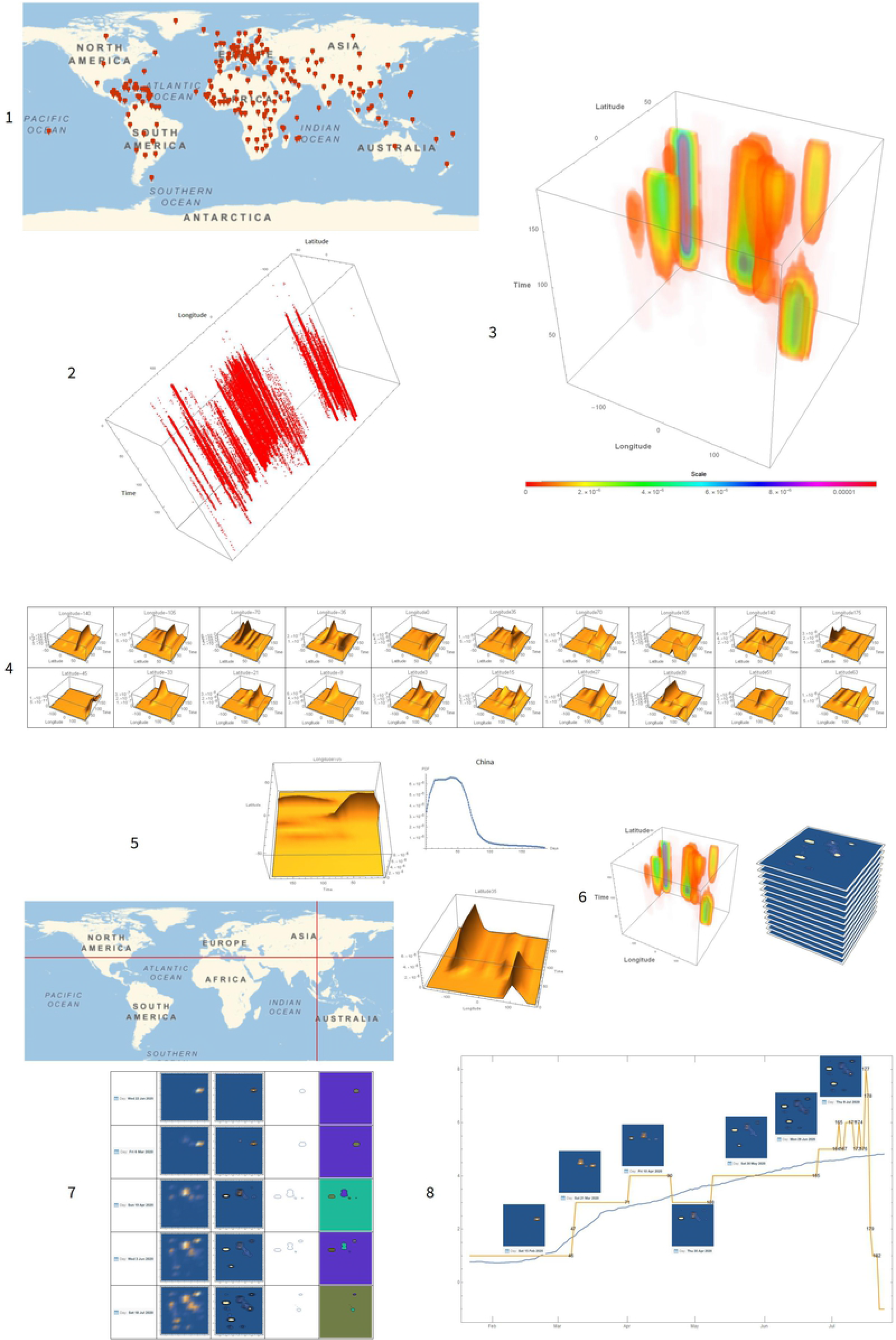
This figure represents the results reported in this work: 1. Geo-referenced data points collection and representation on the Mercator’s Map. 2. Spatial-temporal points distribution obtained by Monte Carlo method simulation. 3. Density plot of the infected probability distribution function (PDF). 4. Spatial representation of the PDF every 20 days. 5. Example of a spatio-temporal representation of the PDF, fixed latitude and longitude. 6. Slicing process of the volume representing the PDF. 7. Detection of morphological components on the slices, obtained in 6. 8. Trends in spatial entropy and the number of morphological components over time.

The considered time runs from January 22, 2020 until July 25, 2020 for a total of 185 days. Obviously, data are not exactly allocated in detailed positions of the space. Each country has had a number of infected individuals, usually collected day by day, but stored as a whole, at the time of the last access to the data. Therefore, to exactly assign data of the infected for each country, a Monte Carlo simulation assigned 2000 points^a^ every day of the considered period, for 370,000, for the 217 nations in the world, according to their relative rate of contagion. Once points have been assigned, they have been randomly scattered on the Mercator’s map, with 2° resolution. This process provided us with a 3D discrete distribution of the Covid-19 infected individuals at the global level ((Fig 1, 2). To attain a good approximation of data with the real virus dynamics in nature, we therefore conjectured both about the spatial density of the randomly distributed points and the relationships between points, as it may allow us to gather information on how contagion has spatially spread and how clusters in identifiable geographical areas have emerged. Accordingly, the spatio-temporal distribution of infected rate has been processed with the Point Process Model (PPM)^12^. To this end, the SKD (Smooth Kernel Distribution) allows to build a distribution function, moving from discrete to continuous, as follows:

**Fig 2.**
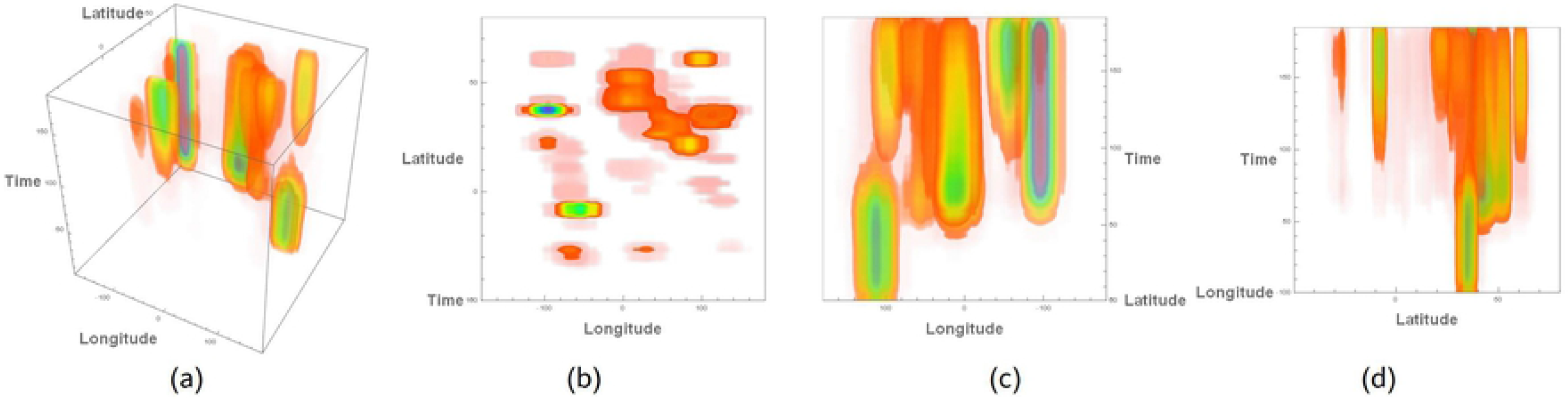
a) 3D Density Plot of the 4D space-time hypersurface describing the probability distribution of rate of Covid-19. b) Orthographic projection from above, with the axes of latitude and longitude. c) Orthographic projection from one side, with the axes of longitude and time. d) Orthographic projection from the other side, with the axes of latitude and time.

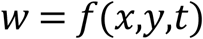

where *x* represents longitude, *y* latitude and *t* the time. This probability distribution is not obtained in closed form, but is proposed through a computational structure in which various interpolating functions are used. The probability distribution function (PDF) *w* = *f*(*x,y,t*) can be built from this distribution. It works as a normal function of three variables. For example, if we want to see how this probability distribution varies in a specific place at a specific time, it is only necessary to provide longitude, latitude and time to get the hypersurface representing the global rates of Covid-19 all over the world (Fig.2).

By observing this hypersurface from different points of view, it is possible to understand the phenomenon of temporal and spatial progression of viral behaviour on a global scale. The first image (Fig. 2a) portrays the final distribution of the virus in space where several clusters are present. In Fig. 2b, the green patterns show the prevalence of contagion in North and South America, with configurations of minor importance. The central pattern is the Eurasian continent, with the Russian one, a little bit detached at the top right. To finish, the red part, at the bottom center, highlights the contagion related to South Africa. The other two images (Fig.2, c and d) allow us to capture the dynamics over time, highlighting the large blocks of contagion that have occurred, especially for the European and Eurasian continents, and, with a little delay, for the American one. However, this representation did not satisfy our expectations, as our main aim was to visualize the virus course as a wave travelling over the world. For this reason, by displaying the 4D hypersurface *w* = *f*(*x,y,t*) and setting a value for *t*, the hypersurface develops in a three-dimensional surface that varies as *t* varies. Consequently, the hypersurface that can only be represented through particular views becomes a surface, as follows:

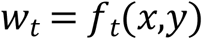

Fig. 3 depicts the grow of these dynamics with a time interval of 20 days, starting from 22 January 2020, every 20 days, for the 185 days considered in this study. The 10 images represent the probability distribution of infected people on a specific day, 20 days apart. Starting from China (January 22), the contagion spread to the Middle East and then Europe (March 2). It is possible to note the progressive growing of the rate of Covid-19 in Europe and the beginning of the pandemic and in North America (April 11). In turn, other geographic areas have been involved with the pandemic spreading, especially Latin America and Russia (May 21), South Africa, the Middle East and India (July 20) (See SI: movie1).

**Fig 3.**
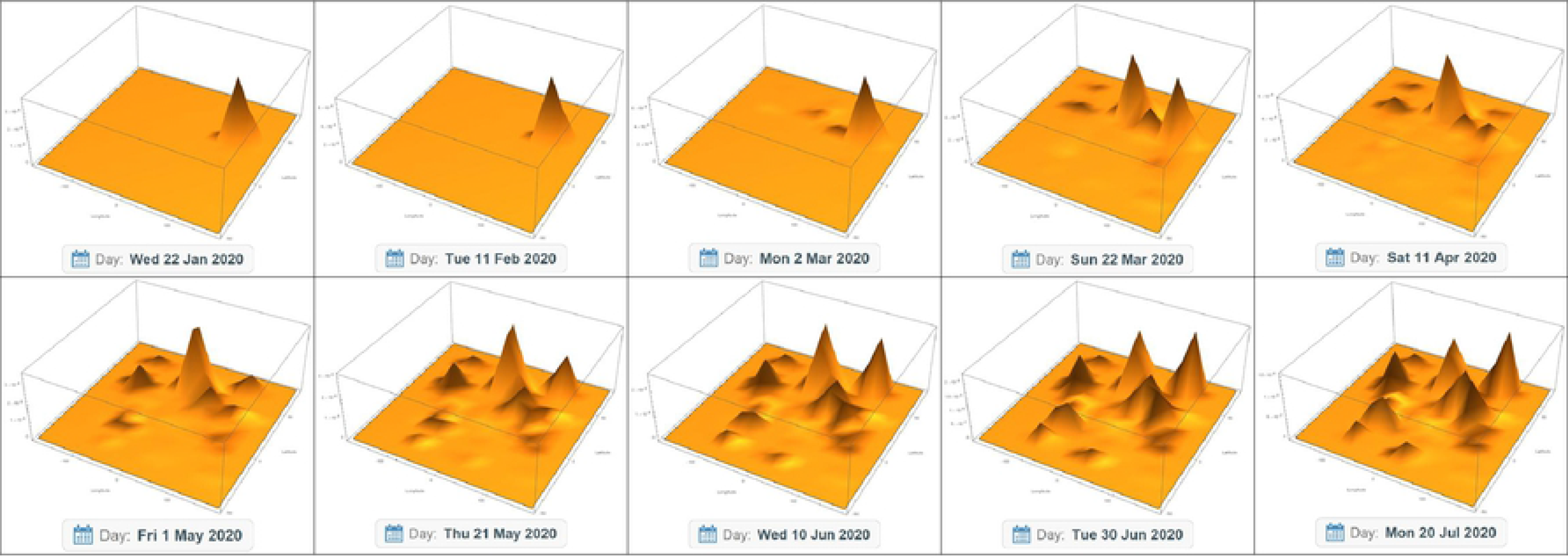
Visualization by a 3D Plot of the 4D space-time hyper surface that describes the probability distribution of rate of Covid-19 with a time window of 20 days, for the 185 days of the pandemic considered in this study.

The method allows visualizing the dynamics of the virus in whatever part of the world. Fig.1, 5 of this work shows and example of this geo-localized analysis of the Covid-19 dynamics. Starting from the 105° est meridian, corresponding to China, it is possible to observe the dynamics of viral growth over time. The presence of the virus in that region is evident. The viral wave continues its course, persisting for a long time in this part of the world and then it almost completely decreases. In this given period, a large part of the world (especially the Pacific area) is essentially free from the pandemic phenomenon, and statistically irrelevant. With the passage of time, however, a wave of increasing importance begins to appear, both in the Russian area and in the two southern parts of India and Central Asia. Visualization on the temporal axis allows, moving from the north along the meridian, to get an idea of the main pandemic development. To enhance further the analysis of the patterning dynamics of the virus in nature, we thought, starting from the four-dimensional structures, to analyse its morphological growth^13^ by considering Covid-19 clusters^14^ formation all over the world. For this purpose, we sliced the volume of the hypersurface along the temporal axes. In analogy with the qualitative changes of chaotic dynamic systems, which realize in the space of the phases the routes to chaos^15^, we have obtained a progression over time of the virus phase changes. By using the Contour Plot function, it is possible to transform three-dimensional maps into two-dimensional maps, essentially detecting contour lines and properly colouring their elevations (Fig.1, 6). By calculating spatial entropy on the slices, quantitative measures of this phenomenon have been obtained. To correlate entropy and the virus phase transitions, we sketched the entropy trend with the curve related to number and size of clusters, detected by the Morphological Component function (Fig.1, 7), emerged during the pandemic process over time. The entropy of the system shows a trend of continuous growth, versus the point of maximum entropy, without highlighting discrete jumps, detected for the transition behaviour of the clusters dynamics evidenced by the morphological components curve (Figure 4). However, on a more accurate analysis, even if the system tends towards a state of maximum entropy, the phase transitions that confirm the evolution of clusters are also noticeable in the entropy curve, in the slopes evidenced by the interpolating lines, as illustrated in the box (top left) of Fig. 4.

**Fig 4.**
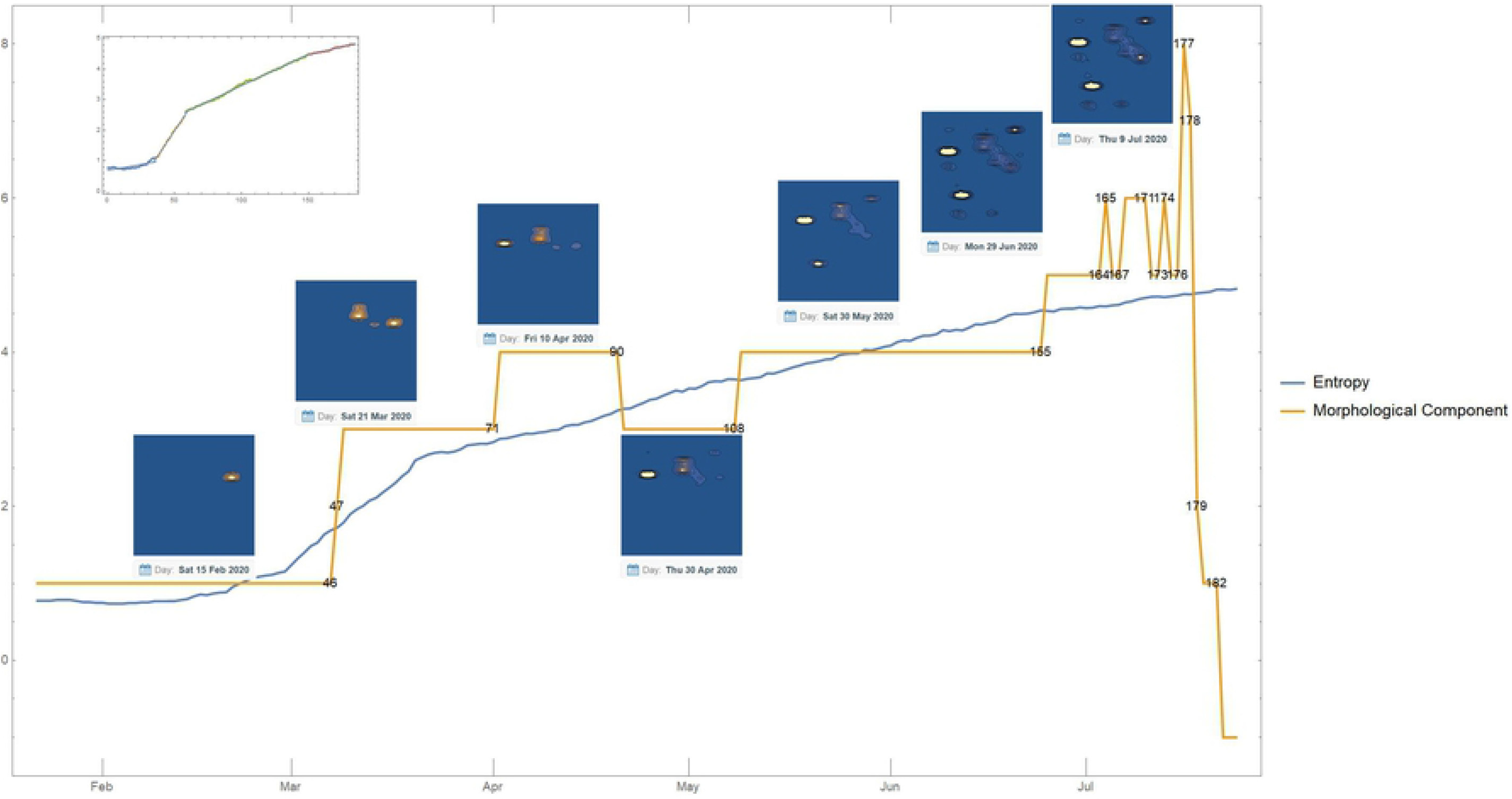
Trends of entropy (blue curve) and cluster number (yellow curve) over time. Blue images represent the emergence of clusters in time. At the top left, the box contains the fitting of curves with the curve of entropy values, calculated on the evolution process of viral patterns.

The numbers on the yellow curve represent the days when a bifurcation occurs, i.e. the emergency, stabilization or rearrangement of pandemic clusters, starting from the day considered as the beginning of the pandemic (22 January 2020), for the 185 days considered in this study. The sequence of numbers in which bifurcations occur is as follows: *Spatio-Temporal_Bifurcations_Days={46,47,71,90,108,155,164,165,167,171,173,174,176,177,178,179,182}*. (See SI: movie2). Results show that, after a variable time of latency, virus bifurcations take place. These bifurcations have temporal rhythms^16^ and occur with a change of place (usually going from one continent to another). Like all biological systems, they seem to have a trend of growth, stability and then decreasing, manifested by the dynamics of clusters. These configurations in turn spread, when they are contiguous in neighbouring continents, then expand enormously, creating completely chaotic zones. This phenomenon occurs after a short period. The system begins to oscillate between different cluster numbers, until it reaches the maximum number of clusters in 177 days, when a collapse of the organized structures occurs. Disintegrations of clusters and an increase in the spatial spread of the virus are observed, as the virus arrives in the US, Brazil, and other countries. Sometimes isolated clusters end, as in the case of China. Other times they maintain an oscillatory behaviour, preserving, however, local minor dynamics that refer to the re-ignition of small outbreaks, in areas where clusters had already reduced. Although there are some similarities between chaotic dynamic behaviour and the dynamics highlighted in this work, identified by the doubling of clusters and the final realization of completely chaotic system configurations, the biological dynamics of the viral spreading seems more variable.

## Methods

The methods developed for this article partially answer these questions:

a. Can virus behaviour be analysed as a complex system?
b. What are the virus main dynamical behaviour?
c. How is it possible to visualize the virus dynamics?
d. Can we highlight spatio-temporal phase transitions, defining both visualization systems and the development of quantitative indicators?

Therefore, we have developed the following steps, most of which developed in Mathematica (https://www.wolfram.com/mathematica/).

### Step 1. Data collection and organization

First, the global collection of geo-referenced data points at a global level, and their representation on the Mercator’s map has been carried out. These points have been acquired from the Wolfram Data Repository^11^, and then connected to geo-spatial references (latitude and longitude for each geographical data point), in order to get data geographically real. Apart from possible errors and variations in the used protocols to collect data, they are very heterogeneous from one country to another. For these reasons, such samples have been cleaned up and considered to be homogenous, overcoming the methodological challenge of using the case-control paradigm and the difficulty of building credible parametric models for human population distributions in a heterogeneous environment^17^. Data from this work were collected until July 25, 2020^18^, for 185 pandemic days starting on January 22, 2020.

### Step 2. Monte Carlo Space-time data plus Probability Distribution Function (PDF)

Monte Carlo simulations have been used to distribute data^19^. Therefore, to allocate rate of infected for each country, a Monte Carlo simulation assigned 2000 points every day of the considered period, for 370,000, for the 217 nations in the world, according to their relative rate of contagion. Once points have been assigned, they have been randomly scattered on the Mercator’s map around the centroid of the nation, in a square of 2° centered on the centroid of the geometric region associated with the nation and, then, visualized for the number of days of the viral course considered in this work (Fig. 1, 2, Main Text). The SmoothKernelDistribution method allows to build a distribution function, moving from discrete to continuous, as follows:

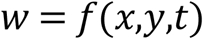

where *x* represents longitude, *y* latitude and *t* time. This probability distribution is not obtained in closed form, but it is used as a computational program in which several interpolating functions are used

The probability distribution (PDF) *w* = *f*(*x,y,t*) can be built from this distribution. It works as a normal function of three variables. For example, if we want to see how this probability distribution varies in specific countries of the world, at a specific time, it is only necessary to provide longitude, latitude and time.

### Step 3. Representing the viral behaviour on a global scale as a hypersurface

The function *w* = *f*(*x,y,t*) is a hypersurface in a four-dimensional space ℝ^4^. Therefore, it cannot be displayed directly. One way to overcome this drawback is to consider *w* as a density associated with each point of ℝ^3^ which overlaps with a particular region of space-time (the spatial part of the region is the geometric representation of the earth in Mercator’s map, while time is the length of the pandemic). Accordingly, the probability density can be represented in every point of the space-time region with a colour code, where the change from red to blue highlights the higher density value of the infected rate probability. The analysis of these images allows us to identify the dynamics of the virus on a large scale (Fig. 1, 3, Main Text).

### Step 4. Reducing the hypersurface for representing the infected probability distribution

A method of visualizing the four-dimensional system *w* = *f*(*x,y,t*) is to set a value of *t*. Thus, the 3D hypersurface becomes a surface, by using the following function:

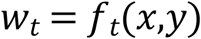

that changes as *t* changes value. We showed some of these views, with time intervals of 20 days from 22 January 2020, for 185 days (Fig. 1, 4, Main Text). A more detailed view of the pandemic course can be seen in the film, which gives day by day the distribution density of infected people globally. This method allow also for looking at the viral by starting by different latitude and longitude of the earth. By choosing longitude and time

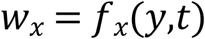

and setting the longitude to different values, it is possible to display the virus probability density distribution on each parallel over time. It is also possible to generate these surfaces by setting the latitude and observing the function over time, as follows:

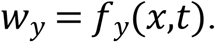

In this way, it is possible to demonstrate that it is possible to visualize these space-time dynamics for each part of the world (Fig. 1, 5, Main Text).

The analysis of these surfaces allows us to interpret the spatial evolution of the virus as a wave, providing a visualization of the pandemic wave that has hit the world.

### Step 5. Slicing process of the hypersurface volume

To further investigate and visualize the emergence of the viral dynamics, considering the virus as a complex system, we thought to cut the volume into slices. A Contour Plot function fulfils this aim transforming three-dimensional maps into two-dimensional maps, essentially detecting contour lines and colouring their elevations appropriately (Fig.1, 6, Main Text). By reducing the hypersurface to slices, it is possible to follow the evolution of the viral dynamics in space-time. The Morphological Components function in Mathematica allows detecting the main structures in these slices. We can consider such structures as attractors, or complex patterns of the virus organization in space and time, that could reside at the *Edge of Chaos*^*20* 21^. They are found where the infected density probability is highest, highlighting patterns, with evolutionary dynamics and morphogenesis processes, identifying phase transitions between order and chaos. Not unlike these systems, the SARS-CoV-2, an RNA virus, needs to replicate and to spread, transmitting and modifying its genetic strand^22^, thus adapting to different ecological niches it is spreading.

### Step 6. Phase transitions detection

The complex dynamics of systems near a phase transition rest on a fundamental capacity for producing information. Therefore, in settings where such phase transitions occur, the virus is capable to generate information. To investigate these processes into quantitative analyses, we thouth to detect the behaviour of the virus on an informational basis. The first approach in this direction is to analyse the entropy of the contagion rate per country. At each interval of time, entropy allows us to evaluate how contagion spreads into the 217 countries considered in this study.

This discreet model does not fully capture the phenomenon for a number of reasons:

a. The number of infected per nation is very different both in spatial size and population, so the number of infected will also be different.
b. The nations are separated and we do not know how they are spatially related to each other.
c. The evidence reported in the previous sections tells us that clusters are transnational and, in fact, neighbouring nations with high levels of infection can be part of the same cluster.

The best way to overcome these criticalities is to use again the PDF that represents a continuous distribution in space and time of the contagion rate. The concept of spatial entropy is closely linked to image analysis and is mainly used in image compression processes^23^. By generating from the PDF the images at each instant of time, it is possible to calculate the entropy associated with each of these images and see how it varies over time. The best way to do this is to build a density plot of the function

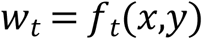

which gives us two-dimensional colour images for each day considered.

The obtained trends are quite close to those of discrete entropy with some notable differences that make this behaviour more likely.

### Step 7. Entropy as a measure of the viral dynamics

The analysis of entropy does not account for the emergence of new attractors. In the complexity approach, the presence of a new attractor (or a new cluster) can be considered a bifurcation point in the pandemic dynamics. One way to evaluate these bifurcation points is to analyse the previous images used in the calculation of entropy to identify the morphological components and to identify when the number of these components changes over time. This kind of analysis gives us the following timescales:

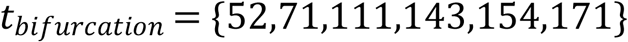

### Step 8. Improving the detection of the viral dynamics

To overcome the reduced sensitivity of the system in intercepting phase transitions of the pandemic system, we adopted an a priori specification of the region that should be properly considered by identifying a minimum probability threshold. Let us consider a minimum probability threshold

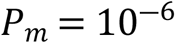

and intersect the probability distributions at each interval of time with a plane with this altitude:

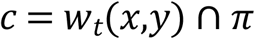

where *t* = 1,2.…185 and π is the plan corresponding to the considered elevation, i.e. it is described by the function *z* = 10^―6^. The result will be a series of closed curves delimiting particular regions of the plan π. The morphological regions will then be exactly these regions. The Density Plot, the ContourPlot together with the intersection curve and the morphological components at the time when only one cluster is detected and at the next time when a new cluster has already formed (Fig. 1. 7, Main Text). The system automatically detects the change of the clusters and therefore the bifurcation points occur at the following times

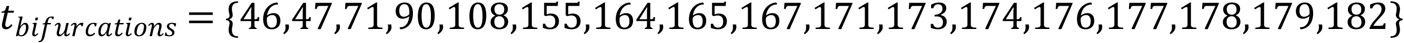

In this way, it is possible to observe how times vary in relation to the entropy curve and the previous calculation of the bifurcations.

The new algorithm seems to represent the real trend of cluster formation and their increased complexity with strong oscillations in the almost final part of the curve. It collapses completely at the end, as the probability distribution becomes so low, due to the decreasing of the default threshold. At these levels of scale, the virus has become so pervasive that its spatial detection is clearly noticeable. The limit of this method concerns the accurate determination of thresholds.

## Conclusions

The scientific meaning of the obtained results can be summarized as follows:

We visualized the patterning dynamics of the novel SARS-CoV-2 virus in every part of the world, considering the virus behaviour as a dynamical complex system, using a 3D Density Plot of the 4D

The space-time dynamics of the virus have been brought to light with the Probability Distribution Function, taken every day from the official beginning of the pandemic for 185 days. This method is very flexible, because since we start from geolocalized data of the infected all over the world, it allows simulating the dynamic trends of the virus in every part of the world. In addition, the other variables of the classic compartmentalized models can be chosen to generate the corresponding dynamics.

The pandemic clusters of infected people have been highlighted, arriving, where data are available, to define even the single infected person in a city street. The geo-spatial clusters of infected people have also been transformed into complex networks, where each node represents an infected person and the links between nodes represent spatial distances. This method allows us to analyse the pandemic phenomenon also through traditional network statistics, establishing the centers of the pandemic in a given region or the diameter of the network, elements that always have a real physical basis.

Finally, the spatio-temporal bifurcations of viral behaviour worldwide have been highlighted, correlating entropy measurements with phase transitions in viral dynamics, as it happens in physical systems.

SI: Movie1 Caption

Dynamic visualization of the trend of the number of infected people worldwide using the Probability Distribution Function (PDF), starting from January 22nd 2020 and ending on July 25th, for a total of 185 days. As the virus spreads, infecting the different continents of the earth, the dynamics become evident in those areas.

SI: Movie2 Caption

Dynamic visualization of the trend of the number of infected people worldwide using the Countour Plot of the Probability Distribution Function (PDF), starting from January 22nd 2020 and ending on July 25th, for a total of 185 days. The movie highlights the patterning phenoma of the viral dynamics all over the world.

## Footnotes

a The choice to use 2000 points at each time step obeys the logic of having the same probability distribution in each time section, in order to compare the distribution in different regions of space, regardless of the number of infected. This allows the analysis of dynamic changes over time of the viral spreading.

## References

1 Wille, M., & Holmes, E. C. The ecology and evolution of influenza viruses. CSH PERSPECT MED, 10 (7), a038489 (2020).

2 Elena, S. F., & Sanjuán, R. RNA viruses as complex adaptive systems. Biosystems, 81(1), 31–41 (2005).

3 Bak P., Tang C., Wiesenfeld K. Scale Invariant Spatial and Temporal Fluctuations in Complex Systems. In: Stanley H.E., Ostrowsky N. (eds) Random Fluctuations and Pattern Growth: Experiments and Models. NATO ASI Series (Series E: Applied Sciences), vol. 157. Springer, Dordrecht (1988).

4 Kitano, H. Computational systems biology. Nature, 420(6912), 206-210 (2002).

5 Chan, J. F.-W. et al.. A familial cluster of pneumonia associated with the 2019 novel coronavirus indicating person-to-person transmission: a study of a family cluster. Lancet 395, 514–523 (2020).

6 Li, R. et al.. Substantial undocumented infection facilitates the rapid dissemination of novel coronavirus (SARS-CoV-2). SCIENCE, 368(6490), 489–493 (2020).

7 Holmes, E. C. The evolution and emergence of RNA viruses. Oxford University Press (2009).

8 Kermack, W. O., & McKendrick, A. G. A contribution to the mathematical theory of epidemics. P R SOC LOND A-CONTA., 115(772), 700–721 (1927).

9 He, S., Peng, Y., & Sun, K. SEIR modeling of the COVID-19 and its dynamics. NONLINEAR DYNAM, 1-14 (2020).

10 Solé, R., & Elena, S. F. Viruses as complex adaptive systems (Vol. 15). Princeton University Press (2018).

11 Wolfram Data Repository on “Epidemic Data for Novel Coronavirus COVID-19” (2020), https://doi.org/10.24097/wolfram.04123.data

12 Wolfram Data Repository on “Epidemic Data for Novel Coronavirus COVID-19” (2020), https://doi.org/10.24097/wolfram.04123.data

13 Kim, M., Paini, D., & Jurdak, R. (2020). Real-world diffusion dynamics based on point process approaches: A review. ARTIF INTELL REV, 53(1), 321–350.

14 Bilotta, E., & Pantano, P. Artificial micro-worlds part II: cellular automata growth dynamics. INT J BIFURCAT CHAOS, 21(03), 619–645 (2011).

15 Bormashenko, E., et al. Clustering and self-organization in small-scale natural and artificial systems. PHIL. TRANS. R. SOC. A, 378(2167), 20190443 (2020).

16 Bilotta, E., Pantano, P., & Stranges, F. A gallery of Chua attractors: Part I. INT J BIFURCAT CHAOS, 17(01), 1–60 (2007).

17 Bertacchini, F., Bilotta, E., & Pantano, P. S. (2020). On the temporal spreading of the SARSCoV-2. Preprint at medRxiv. https://www.medrxiv.org/content/10.1101/2020.08.01.20166447v1 (2020).

18 Diggle, P. J. Statistical analysis of spatial and spatio-temporal point patterns. CRC press (2013).

19 WHO, report 187; https://www.who.int/docs/default-source/coronaviruse/situationreports/20200725-covid-19-sitrep-187.pdf?sfvrsn=1ede1410_2.

20 Baddeley, A. et al.. Residual analysis for spatial point processes (with discussion). J ROY STAT SOC B, 67(5), 617–666 (2005).

21 Langton, C. Computation at the edge of chaos: Phase transition and emergent computation (No. LA-UR-90-379; CONF-8905201-5). Los Alamos National Lab., NM (USA) (1990).

22 Sun, G. Q. et al. Pattern transitions in spatial epidemics: Mechanisms and emergent properties, PHYS. LIFE REV., 19, 43–73 (2016).

23 Forster, P., Forster, L., Renfrew, C., & Forster, M. Phylogenetic network analysis of SARS-CoV-2 genomes. PROC. NATL. ACAD. SCI. USA, 117(17), 9241–9243 (2020).

24 Dhawan, S. A review of image compression and comparison of its algorithms. AEU-INT J ELECTRON C, 2(1), 22–26 (2011).

